# Age-related fornix decline predicts conservative response strategy-based slowing in perceptual decision-making

**DOI:** 10.1101/2023.11.15.567204

**Authors:** Lauren Revie, Claudia Metzler-Baddeley

**Author notes:** Corresponding author: Claudia Metzler-Baddeley.

## Abstract

Aging leads to increased response latencies but the underpinning cognitive and neural mechanisms remain elusive. We modelled older and younger adults’ response time (RT) data from a 2-choice flanker task with a diffusion drift model (DDM) and employed multi-shell diffusion weighted magnetic resonance imaging and spectroscopy to study neurobiological predictors of DDM components thought to govern RTs: drift rate, boundary separation and non-decision time. Microstructural indices of fractional anisotropy (FA), diffusivities and the restricted signal fraction (FR) from the Composite Hindered and Restricted Model of Diffusion (CHARMED) were derived from white matter pathways of visuo-perceptual and attention networks (optic radiation, inferior and superior longitudinal fasciculi, fornix) and estimates of metabolite concentrations [N-acetyl aspartate (NAA), glutamate (Glx), γ-aminobutyric acid (GABA), creatine (Cr), choline (Cho) and myoinositol (mI)] were measured from occipital (OCC), anterior and posterior cingulate cortices (ACC, PPC). Ageing was associated with increased RT, boundary separation, and non-decision time. Differences in boundary separation but not non-decision time mediated age-related response slowing. Regression analyses revealed a network of brain regions involved in top-down (fornix FA, diffusivities in right SLF) and bottom-up processing (mI in OCC, AD in left optic radiation) and verbal intelligence as significant predictors of RTs and non-decision time (NAA in ACC, AD in the right ILF, creatine in the OCC) while fornix FA was the only predictor for boundary separation. Fornix FA mediated the effects of age on RTs but not *vice versa*. These results provide novel insights into the cognitive and neural underpinnings of age-related slowing.

## 1. Introduction

One of the best-established findings in aging research concerns the slowing of response speed (Salthouse, 1979) that can be observed in a wide range of tasks involving simple decision-making to more complex executive functioning with spatial-perceptual discrimination being disproportionally affected (Ratcliff et al 2007; Verhaegen & Cerella. 2008).

Age-related slowing is often accompanied by a lengthening of the speed accuracy trade-off (SAT) which refers to the trade-off that occurs between responding as timely and as accurately as possible when completing a time-pressured cognitive task. Greater SATs in aging are thought to occur due to older adults adopting a more cautious response strategy that favours response accuracy over speed (Starns & Ratcliff, 2010). In contrast, younger adults typically make faster responses which may be at greater risk of errors (Der & Deary, 2005). It is commonly thought that the change to a more cautious response strategy at the cost of slower reaction times (RT) arises due to an age-related decline in sensorimotor functions leading to a distrust in being able to provide a correct response.

Impairments in sensorimotor functions may affect both bottom-up sensory and top-down decision-making and motor execution processes (Singh et al., 2013). According to the sensory degradation hypothesis (Hurley et al., 1998; Zalewski, 2015), age-related deterioration in sensory functions results in noisier sensory input and hence longer perceptual processing time for effectively interpreting a stimulus. This in turn increases overall RTs or the likelihood of an incorrect response if RT is not lengthened (Basak & Verhaeghen, 2011). Indeed, age-related perceptual decline has been observed at some of the lowest levels of visual processing such as visual contrast, even in those with intact visual acuity and an absence of visual impairment (Delahunt et al., 2008, Elliott et al., 1990, Govenlock et al., 2009).

In addition, the “slowed motor response” hypothesis proposes that age-related increases in RTs emerge from a slowing of top-down decision-making and motor generation and execution processes (Bashore et al, 2015; Falkenstein et al 2006). This view is backed up by evidence from electroencephalogram studies suggesting that older people are most compromised at the interface of translating a stimulus input into a response output (Bashore et al, 2015; Kubo-Kawai & Kawai, 2010).

It is plausible that both sensory degradation and motor noise contribute to a slowing of decision-making processes in ageing. However, the analysis of overall RTs or SATs alone, does not allow for a separation of the distinct cognitive components that may contribute to age-related slowing.

Sequential-sampling models (Heitz, 2014; Stafford et al 2020), such as Ratcliff’s drift diffusion model (DDM) can be applied to RT data from choice RT tasks to estimate parameters that map distinct cognitive components involved in decision making. The DDM approach has been employed by several studies to clarify the cognitive underpinnings of older adults’ slowed RTs and SATs (see for review Theissen et al 2021).

According to the DDM, sensory input provides information that accumulates over time. This information fluctuates randomly between two thresholds: a lower threshold representing an incorrect response choice and an upper threshold representing the correct response choice (Figure 1). When the accumulated information, after some time, crosses one of these thresholds, it triggers the corresponding response. The main components of the DDM, that control the time it takes to reach one of these thresholds, i.e., the response time, include the speed of information uptake (drift rate) (Figure 1 blue), the distance between the thresholds, that reflects the degree of conservatism regarding the response criterion (boundary separation) (Figure 1 green), and the time required for non-decisional processes including sensory-perceptual encoding and motor response execution (non-decision time) (Figure 1 red). If the boundary separation decreases (while keeping drift rate non-decision time constant), the response time becomes shorter, but the likelihood of making an error increases. Conversely, if the boundary separation increases (again with drift rate and non-decision time held constant), the response times lengthens, but the likelihood of making an error decreases. Thus, according to Ratcliff’s DDM model, the distance between the thresholds, i.e., the boundary separation value, determines the length of the SAT.

**Figure 1.**
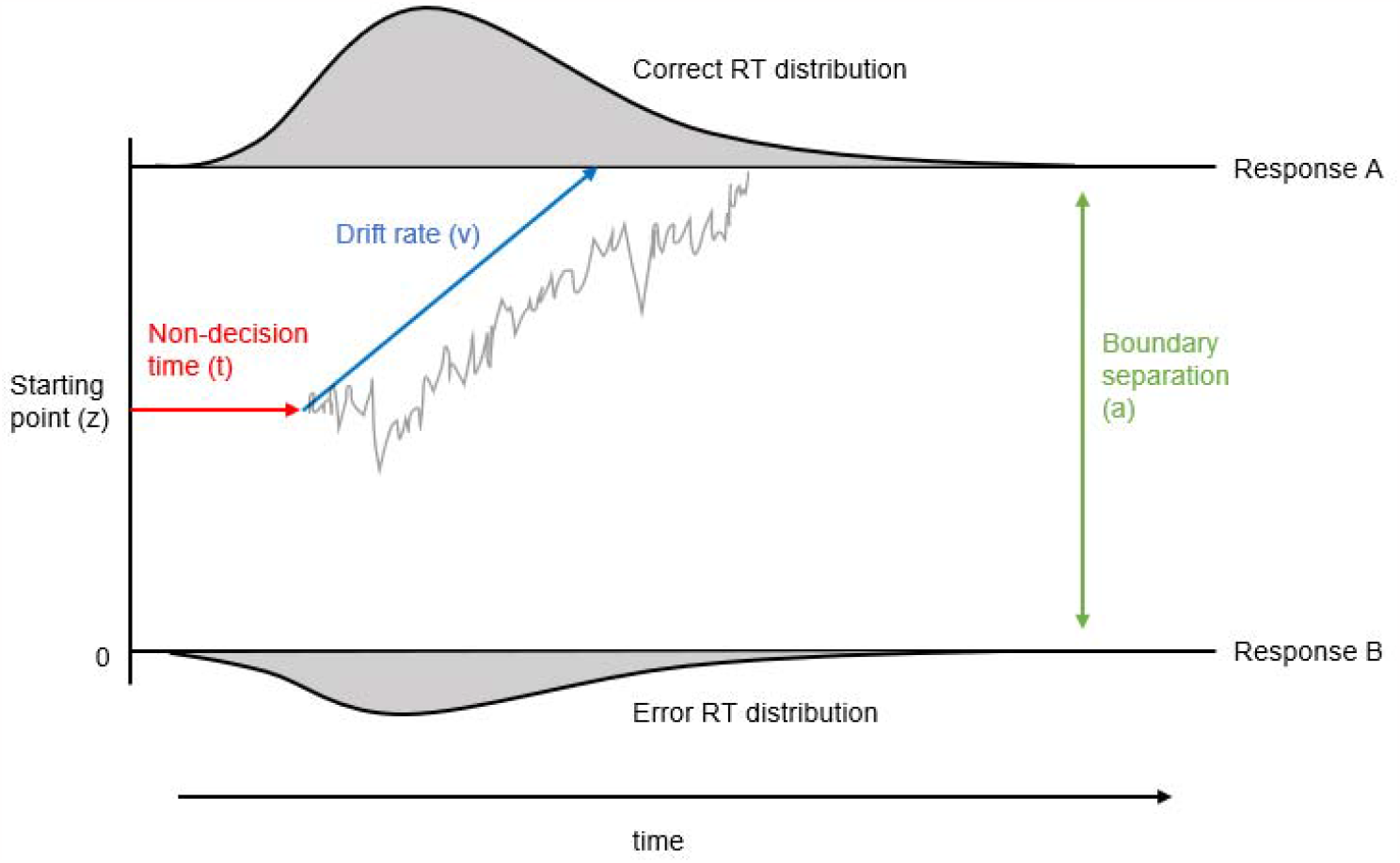
The drift diffusion model (DDM) of response time. UppeThese results provide novel insightsr and lower lines ‘Response A’ (correct) and ‘Response B’ (incorrect) denote the different responses in a 2-choice task (for example, left or right key press). Reac-tion times (RT) are fit to the DDM to return an estimate of non-decision time (t) (red) reflecting perceptual and motor processing time, boundary separation value (a) (green), reflecting the distance between the two response criterion thresholds for A or B that reflects the amount of information that needs to be accumulated to trigger a response, and drift rate (v) (blue) reflecting the efficiency of the drift process. Information is thereby assumed to accumulate in a random walk-like diffusion process (grey wiggly line) that commences at the starting point toward one of the two response boundaries. Adapted from Kühn et al (2011).

Indeed, in the literature higher boundary separation values and longer non-decision times have been reported consistently in older compared with younger individuals while differences in drift rates were found to be moderated by task type (e.g. episodic versus semantic memory) and difficulty (Theisen et al 2021; Rabbitt, 1979).

To date only a few studies have investigated the neurobiological underpinnings of age-related differences in DDM parameters (Forstmann et al., 2011, Kühn et al., 2011; Madden et al., 2020,;Monge et al., 2017, Yang et al., 2015). Magnetic resonance imaging (MRI) studies have found age-related increases in boundary separation to be associated with reduced striatal activity (as measured with the blood oxygen level dependent (BOLD) signal) (Kühn et al., 2011) and with reduction in fractional anisotropy (a diffusion tensor imaging (DTI) based measurement of fiber directionality/coherence) in white matter connections between the striatum and the pre-Supplementary Motor Area (preSMA) (Forstman et al (2011). Other studies have linked age-related increases in non-decision time (Madden et al., 2010; Madden et al 2020) and in drift rate (Meerestein, Mullin & Madden, 2023) to differences in the BOLD signal in frontoparietal regions. These findings are consistent with evidence suggesting that reductions in fronto-parietal activity may underpin age-related slowing in simple and choice RT tasks and in SAT (Bugg et al., 2006, Jackson et al., 2012, Madden et al., 2004, Monge et al., 2017). They also accord with well-documented evidence of structural and functional changes in fronto-parietal and striatal networks with age (Olesen et al., 2003; Raz et al., 2003) that are thought to be important for attention control and decision making (Macpherson et al., 2014; Markett et al., 2015). In summary, the findings from DDM based studies suggest that age-related increases in RTs and SAT may be driven by non-decision related sensorimotor decline (non-decision time) and by longer processing times required before a decision threshold can be reached (boundary separation). They further suggest that age-related neural decline in fronto-parietal and striatal decision-making networks contribute to the differences in DDM components in ageing. However, the precise cognitive and neurobiological mechanisms that underpin age-related increases in DDM parameters and response slowing remain unknown.

The aim of the present study was to further elucidate the cognitive and neurobiological substrates of age-related slowing in visual-perceptual discrimination using an Eriksen flanker choice test (Eriksen & Eriksen, 1974). The Eriksen flanker task is a classic response inhibition task that involves the presentation of a target arrow flanked by distractor arrows, which are either congruent with the directional response to the target, i.e., a left or right key press, incongruent (pointing into the opposite direction), or neutral. Younger (n = 25, age range = 18-29 years) and older participants’ (N = 25, age range = 62-80 years) RT data from the flanker tasks were modelled to derive SAT and DDM parameters. Correlation coefficients between RTs, SAT, and DDM-derived parameters were calculated and mediation analysis (Hays et al., 2012) was used to identify those DDM component(s) that accounted for the shared variation in RTs and SAT across both groups. Gray matter metabolic and white matter microstructural measurements were derived from key regions of interest (ROIs) within visual-perceptual and decision-making networks and hierarchical regression analyses were conducted to identify neurobiological brain predictors of cognitive components.

For this purpose, we employed multi-shell high angular resolution diffusion imaging (msHARDI) (Descoteaux, 1999) to quantify white matter microstructural properties and magnetic resonance spectroscopy (MRS) to measure gray matter metabolite concentrations. These modalities were chosen because it is well established that microstructural properties of white matter brain connections, that allow the efficient communication within and between brain networks, deteriorate with advancing age and contribute significantly to cognitive decline including response slowing in ageing (Kuznetzova et al 2016). Most studies that have investigated age effects on white matter microstructure employed diffusion tensor imaging (DTI) and have consistently found reduced fractional anisotropy (FA) and increased mean diffusivity (MD), axial diffusivity (AD) and radial diffusivity (RD) in older relative to younger participants (Burzynska et al., 2010; de Groot et al., 2015). Age-related decline of white matter microstructure occurs across the whole brain but is particularly apparent in fronto-parietal and limbic regions (Charlton et al., 2010) and correlates with age-related differences in processing speed, episodic memory, and executive functions (Kerchner et al., 2012; Kuznetsova et al., 2016).

Here we studied the microstructure of the following white matter pathways that connect visual and attention network regions and are known to be involved in top-down and/or bottom-up visual perceptual processing and attentional functioning: the optic radiation, that connects the lateral geniculate nucleus with the primary visual cortex in the occipital lobe and is important for bottom-up visual sensory processing (Schurz et al., 2014); the inferior longitudinal fasciculus (ILF), that connects occipital and anterior temporal cortices and is involved in bottom-up visual-perceptual object, face, and place processing; the fornix, the main output tract of the hippocampus to other limbic and cortical regions, that mediates mnemonic and complex visual discrimination functions (Lech et al 2016); and the superior longitudinal fasciculus (SLF), that connects parietal with prefrontal cortices, notably the right SLF being the crucial white matter pathway of the top-down right-lateralized attention-executive network (De Schotten et al., 2011). White matter microstructural properties of these tracts were not only characterised with DTI metrics (FA, MD, RD, AD) but also with the restricted signal fraction FR, a proxy index for axonal density, from the Composite Hindered and Restricted Model of Diffusion (CHARMED) (Assaf & Basser, 2004). FR is thought to be a valuable metric to quantify in this context, as it has been shown to be more sensitive than DTI indices and has been suggested as a potential biomarker for axonal microstructure changes (De Santis et al., 2017) which are well-established in aging.

Furthermore, ageing is also known to be associated with changes in concentrations of metabolites that are important for healthy neuronal functioning. More specifically, older compared with younger adults show reduced concentrations of N-acetyl aspartate (NAA) (Lu et al., 2004), an estimate of neuronal density and function, and of glutamate/glutamine (Glx) and γ-aminobutyric acid (GABA), the major excitatory and inhibitory neurotransmitters in the brain ((Rae et al., 2014; Stagg & Rothman, 2013). In addition, ageing has been found to be associated with increased concentrations of creatine (Cr), choline (Cho) and myoinositol (mI), that have been linked to inflammation, demyelination, and glia cell proliferation (Glanville et al., 1989; Zeisel & da Costa, 2009). Such age-related related differences in metabolites have been observed in many brain regions including the occipital, frontal, and anterior and posterior cingulate cortices (Chiu et al., 2014; Gruber et al., 2008; Haga et al., 2009; Gao et al., 2013; Marjańska et al., 2017; Pitchaimuthu et al., 2017; Porges et al., 2017; Simmonite et al., 2019). Here we measured concentrations of NAA, Glx, GABA, creatine, choline and mI in the following three cortical ROIs: the occipital cortex (OCC), the anterior cingulate cortex (ACC) and the posterior parietal cortex (PPC). These ROIs were selected as they form key regions of visual perception and attention networks that mediate bottom-up and top-down processing streams with OCC being involved in primary visual processing (Pitchaimuthu et al., 2017), PPC mediating sensory-perceptual integration (Chiu et al., 2014) and ACC playing a key-role in decision-making by means of error signalling and event rewarding (Weerasekera et al., 2020).

In this way we were able to characterise age-related metabolic and microstructural differences in key structures of the visual and attentional networks and assess whether these brain differences were predictive of differences in RT, SAT, and DDM parameters. Based on above summarized findings, we hypothesized that aging would be associated with increases in RT, SAT, boundary separation, and non-decision time as well as with reductions in NAA, GABA, Glx, FA and FR and increases in choline, myoinositol, glutamate, MD, RD and AD in all grey matter ROIs and white matter pathways. We further hypothesized, that age-related metabolic and microstructural differences in both bottom-up sensory processing areas (OCC, optic radiation, ILF) and top-down motor execution areas (SLF, fornix, ACC, PPC) would account for differences in overall RTs, and in non-decision time, as the latter reflects both sensory and motor execution processes. In contrast, only regions involved in top-down decision-making (SLF, fornix, ACC, PPC) were expected to predict differences in SAT and boundary separation. No specific hypotheses regarding drift rate were generated given the ambiguity of findings in the literature.

## 1. Methods

### 2.1 Participants

Participants were recruited from the School of Psychology community participant panel at Cardiff University and consisted of younger (aged 18-29) and older (aged 62-80) adults. Twenty-five participants were recruited into each group, all of whom provided informed written consent prior to taking part in the study in accordance with the Declaration of Helsinki (Cardiff University School of Psychology Ethics committee reference 18.06.12.5313; NHS Research Ethics committee REC reference 18/WA/0153). All participants were cognitively healthy, i.e., had a Montreal Cognitive Assessment (MOCA) score ≥ 26. Participants also completed MRI screening prior to the study, excluding any participants with MRI contraindications such as metallic or electronic bodily implants, some dental work and some tattoos, subject to radiographer assessment. Individuals with visual impairments, such as visual field loss or glaucoma were also excluded from the study. Table 1 summarises participants’ demographic information as well as their mean performance on cognitive and visual screening tasks. Both groups were comparable with regards to sex, handedness, years of education, and visual acuity. All participants had normal or corrected normal visual acuity with Snellen Fractions ≥ 1. The young group performed slightly better on the Test of premorbid functioning UK version (TOPF-UK) (Wechsler et al., 2011), which involves reading out a list of irregular words, and provides an estimate of verbal intelligence.

**Table 1.** Demographic and baseline cognitive scores for younger and older adults. Mean and standard deviation (SD) for younger and older adults’ performance. MOCA = Montreal cognitive assessment, TOPF-UK = test of premorbid functioning, UK-edition.

|  | Younger Mean (SD) | Older Mean (SD) | t<sub>(49)</sub>-value<br>(p-value) |
| --- | --- | --- | --- |
| Age | 21.56 (2.76) | 68.36 (6.1) | 34.8 (< 0.001) |
| Sex | Male (8) Female (17) | Male (12) Female (13) | - |
| Handedness | Left (1) Right (24) | Left (4) Right (21) | - |
| Years of education | 16.08 (2.3) | 15.64 (4.3) | 0.45 (0.66) |
| MOCA score | 28.84 (1.25) | 29.04 (1.1) | 0.58 (0.56) |
| Visual acuity (Snellen fraction) | 1.91 (0.18) | 1.77 (0.3) | 1.8 (0.08) |
| TOPF-UK score | 60 (6.43) | 64.72 (4.5) | 3.0 (0.004) |

### 2.2 Materials & Procedure

#### 2.2.1 Cognitive and visual testing

Testing was conducted at Cardiff University Brain Research Imaging Centre (CUBRIC), during one visit lasting approximately 2 hours. Participants completed a visual acuity task and flanker task on a computer which is described in detail below. The task was presented on a 15” screen (1440 x 900 native resolution) and responses were recorded using a wireless keypad. The flanker task was written by LR using PsychoPy psychophysics software (Peirce, 2009) for Python (v1.85.6) following the original methodology of the Attention Network Task (ANT) (Fan et al., 2002) unless otherwise stated. Visual acuity was assessed using the Freiburg Visual Acuity and Contrast Test (FRACT; Bach, 1996). Participants viewed the screen from a distance of 2m (as recommended by test manufacturers) and responded to circular stimuli, where the target was a ‘gap’ in the circle. Stimuli was reduced in size for each correct trial to achieve a Snellen fraction measure of visual acuity.

Reaction times (RTs) were recorded using a modified Attention Network Test (ANT) flanker task (Fan et al., 2002), and speed accuracy trade-off and DDM parameters were calculated using these RTs. The modified ANT stimuli consisted of five horizontal arrows presented on the screen in which participants were instructed to attend to the central arrow as the target. Central arrows were flanked by horizontal lines (neutral condition), arrows facing in different directions to the target (incongruent condition), or arrows facing in the same direction as the target (congruent condition). During this version of the ANT, stimuli were presented in the same central position on the screen following the presentation of a fixation cross. Participants viewed the screen from a seated position, 400mm from the computer screen. Participants were instructed to maintain focus on the central fixation point of the screen and respond as quickly and accurately as possible. In accordance with the original study (Fan et al., 2002), these stimuli subtended 3.08° of visual angle. Fixations were presented for a random variable length of time between 400-1600ms and target stimuli were presented for a maximum of 1700ms. Participants completed 96 trials (32 trials per condition) in each block, for a duration of 5 blocks. Between blocks, participants were instructed to rest for 30 seconds, before being given a 5 second count-down into the following block. The entire task totalled 480 trials and took approximately 12-15 minutes to complete.

#### 2.2.2 Speed accuracy trade-off (SAT) calculation

Speed accuracy trade-off (SAT) was calculated from RTs using the linear integrated speed accuracy score (LISAS; Vandierendonck, 2017) method, which combines RT and proportion of error in a linear manner, according to the formula (Equation 1).

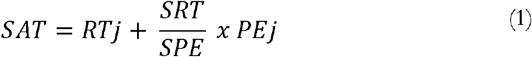

Where RTj is the mean RT, PEj is the proportion of errors, SRT is the participants’ overall RT standard deviation, and SPE is the participants’ overall standard deviation for the proportion of errors. To assess correlations between RT, SAT, and DDM indices, Spearman’s Rho correlation coefficients were calculated between these measurements. Linear mediation analysis was then used to test for the indirect effects of DDM mediator variables on the direct effects of SAT on mean RT. The significance of indirect and direct effects was assessed with a 95% confidence interval based on bootstrapping with 5000 replacements.

#### 2.2.3 Drift diffusion modelling (DDM)

DDM parameters were calculated using the EZ DDM model (Wagemakers et al., 2007) which was incorporated into an in-house R based custom script. Raw RT and accuracy data for each participant for congruent, neutral and incongruent trial conditions were input into the script in R Studio (v 1.1.463). The script first calculated means and variances of correct RTs. Incorrect trials were not included in the remainder of the analysis (average retained trials = 469). Following this, the script calculated DDM parameters using the equation provided in Wagenmakers et al., (2007) under the assumption that trial-to-trial variability was zero and the starting point of each decision process was equidistant from the response boundaries (Schmiedek et al., 2007). This resulted in average estimates for non-decision, boundary separation and drift rate for each participant. Details of the mathematical basis for the EZ model can be found in Wagenmakers, Van der Maas & Grasman (2007).

#### 2.2.2 Magnetic Resonance Imaging (MR) Imaging and Spectroscopy

##### 2.2.2.1 MR data acquisition

All MR data were acquired on a Siemens 3 Tesla (T) Magnetom Prisma MR system (Siemens Healthcare GmbH, Erlangen) fitted with a 32-channel receiver head coil at CUBRIC. A 3D, T1-weighted magnetization prepared rapid gradient-echo (MP-RAGE) structural scan was acquired for each participant (TE/TR = 3.06/2250ms, TI = 850ms, flip angle = 9deg, FOV = 256mm, 1 x 1 x 1mm resolution, acquisition time = ∼6min). The MPRAGE was used as anatomical reference for the placement of MRS region of interest voxels.

MRS was used to acquire frequency spectra to quantify metabolites of Glx, GABA, NAA, choline, creatine and myoinositol. Single voxel proton spectra were obtained from voxels of interest placed in the occipital cortex (OCC, voxel measuring 30 x 30 x30 mm^3^), the posterior parietal cortex (PPC, voxel measuring 30 x 30 x 30 mm^3^) and the anterior cingulate cortex (ACC, voxel measuring 27 x 30 x 45mm). The OCC voxel was placed above the tentorium cerebelli, avoiding scalp tissue in order to prevent lipid contamination to the spectra. The PPC voxel was placed with the posterior edge against the parieto-occipital sulcus, and the ventral edge of the voxel above and parallel to the splenium. Finally, the ACC was placed directly dorsal and parallel to the genu of the corpus callosum. In each voxel, a spectral editing acquisition (MEGA-PRESS, Mescher et al., 1998) was performed, involving applying an additional pulse symmetrically about water resonance, providing ‘on’ and ‘off’ editing pulses which allow for the subtraction of peaks which may mask GABA in the spectra (TE/TR = 68/2000ms, 168 averages, acquisition time = ∼12 min per voxel). Manual shimming was performed before all MRS scans to ensure water-line width of 20Hz or lower, in order to obtain accurate peaks in the spectra (Figure 2).

**Figure 2.**
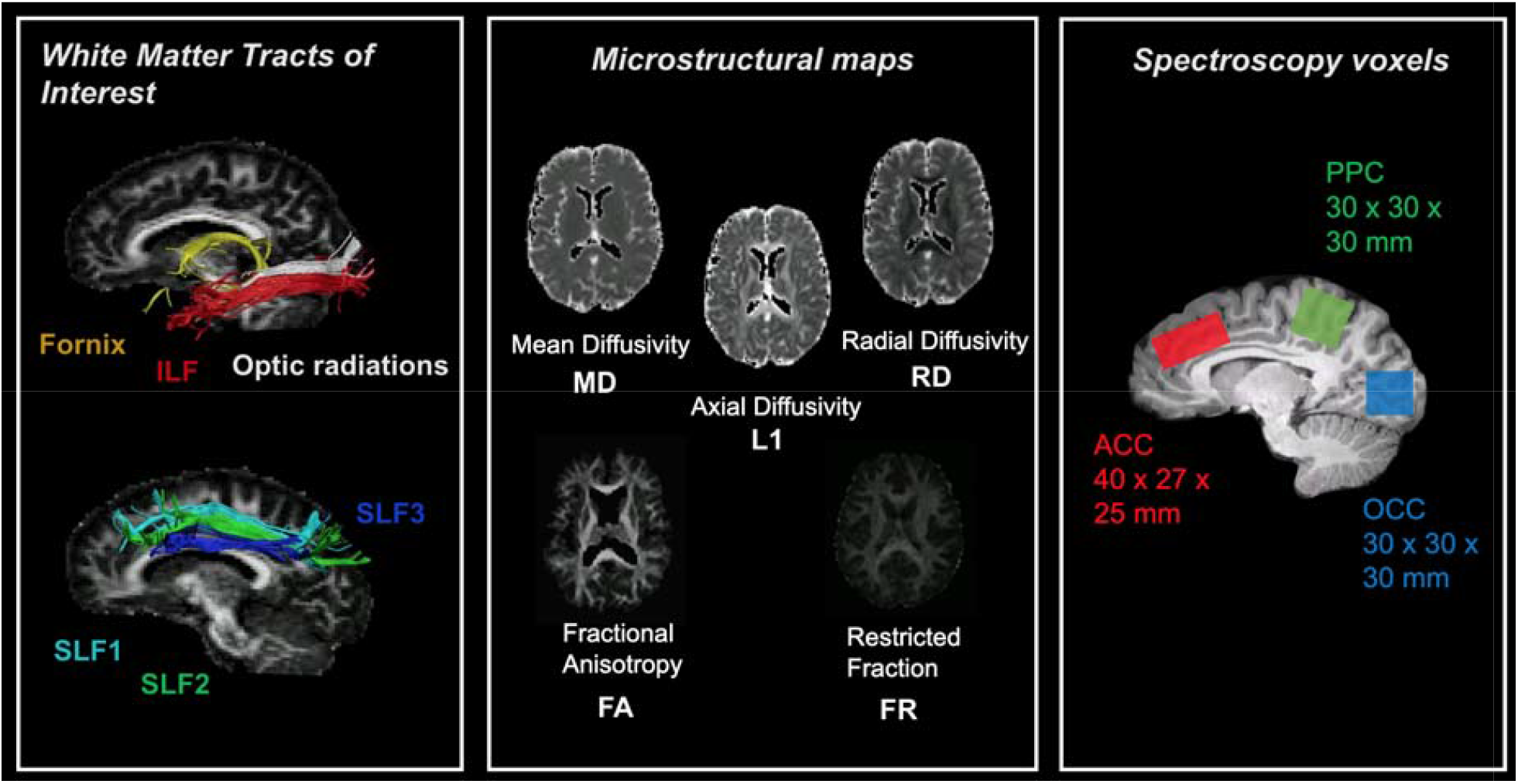
White matter tracts, microstructural maps and location of spectroscopy voxels for measurements of interest. ACC = anterior cingulate cortex, ILF= inferior longitudinal fasciculus, PPC = posterior parietal cortex, OCC= occipital cortex, SLF = superior longitudinal fasciculus.

A multi-shell diffusion MRI sequence was also conducted using a high angular resolution diffusion (HARDI) weighted echo-planar imaging (EPI) sequence (TE/TR = 73/ 4100ms, FOV = 220×220mm, isotropic voxel size 2mm^3^, 66 slices, slice thickness 2mm, acquisition time ∼15 min, 2 × 2 × 2mm resolution). Five diffusion weightings were applied along gradient directions: b = 200 s/mm^2^ (20 directions), b= 500 s/mm^2^ (20 directions) b = 1200 s/mm^2^ (30 directions), b=2400 s/mm^2^ (61 directions), b =4000 s/mm^2^ (61 directions). 12 unweighted (b0) volumes were acquired, interspersed throughout diffusion-weighted scans. In addition, a diffusion reference sequence was acquired for later blip-up blip-down analysis to correct for EPI distortion (Bodammer et al., 2004) in which a diffusion weighting of b=1200 s/mm^2^, and 12 un-weighted (b0) images were acquired interspersed throughout the sequence (Figure 1). Multi-shell diffusion weighted imaging data were acquired to fit the diffusion tensor and the Composite Hindered and Restricted Model of diffusion (CHARMED) (Assaf & Basser, 2005) to gain microstructural maps of fractional anisotropy (FA), mean diffusivity (MD), radial diffusivity (RD), axial diffusivity (L1), and restricted signal fraction (FR).

##### 2.2.2.2 MR analysis

MRS data were analysed using Totally Automatic Robust Quantification in NMR (TARQUIN) version 4.3.11 (Reynolds, Wilson, Peet & Arvanitis, 2006) in order to determine estimated concentrations of other metabolites of interest (Choline, NAA, Glx, Creatine, Myoinositol). To ensure data quality, metabolites were excluded if the Cramer Rao Lower Bound (CRLB) was above 20% as recommended (Stagg & Rothman, 2013). MEGA-PRESS data were analysed using GANNET (GABA-MRS Analysis Tool) version 3.0 (Edden et al., 2014). Estimated metabolite values were corrected to account for cerebrospinal fluid (CSF) voxel fraction, and water reference signal was corrected to account for differing water content of CSF, grey matter and white matter. All metabolites were quantified using water as a concentration reference and were expressed as concentration in millimoles per unit (mM).

Two-shell HARDI data were split by b-value (b=1200, and b=2400 s/mm^2^) and were corrected for distortions and artifacts using a custom in-house pipeline in MATLAB and Explore DTI (Leemans et al., 2009). Correction for echo planar imaging distortions was carried out by using interleaved blip-up, blip-down images. Tensor fitting was conducted on the b=1200 s/mm^2^ data, and the two compartment ‘free water elimination’ (FWE) procedure was applied to improve reconstruction of white matter tracks close to ventricles (Pasternak et al., 2009) and to account for partial volume contamination due to CSF which is particularly apparent in older age (Metzler-Baddeley et al., 2012). Data were fit to the CHARMED model (Assaf & Basser, 2005) which involved the correction of motion and distortion artefacts with the extrapolation method of Ben-Amitay et al., (2012). The number of distinct fibre populations in each voxel (1, 2 or 3) was determined using a model selection approach (De Santis et al., 2014) and FR maps (Assaf & Basser, 2005) were then extracted by fitting the CHARMED model to the DWI data, with an in-house script. This resulted in FA, MD, RD, AD and FR maps.

Whole brain tractography was then performed with the dampened Richardson-Lucy (dRL) spherical deconvolution method (Dell’Acqua et al., 2010). Tractography was performed on the b=2400 s/mm^2^ data to provide better estimation of fibre orientation (Vettel et al., 2012). The dRL algorithm extracted peaks in the fibre orientation density function (fODF) in each voxel using a step size of 0.5mm. Streamlines were terminated if directionality of the path changed by more than 45 degrees using standardised inhouse processing pipeline at CUBRIC. Manual fibre reconstructions were performed in ExploreDTI v4.8.3 (Leemans et al., 2009). Tracts of interest were manually drawn on direction encoded colour FA maps in native space. ILF reconstruction was obtained according to protocol by Hodgetts et al. (2015) and Wakana et al. (2007). The SLF was subdivided into three subdivisions, the SLF1, 2 and 3 which were delineated according to protocol by De Schotten et al., <u>(2011)</u>. The SLF was subdivided based on the distinct contributions of each tract to different functions of attention and executive processing; the SLF 1 being associated with spatial functions and goal-directed attention (De Schotten et al., 2011; Parlatini et al., 2017), the SLF2 being associated with orienting attention and integration of dorsal and ventral attention networks (Nakajima et al., 2019), and the SLF 3 being associated with reorienting of spatial attention (De Schotten et al., 2011; Nakajima et al., 2019). The fornix was reconstructed by locating the body of the fornix bundle according to Metzler-Baddeley et al. (2011), and the optic radiation was delineated by placing a seed region on the white matter of the optic radiation lateral to the lateral geniculate nucleus in the axial plane (Thompson et al., 2014) (Figure 2).

#### 2.2.2 Statistical Analysis

Statistical analyses were conducted in R-studio (v 1.1.463), SPSS version 27 (IBM) and the PROCESS computational tool for mediation analysis version 4.3 (Hayes, 2012). Data were assessed for normality with the Kolmogorov-Smirnov test and were either analyzed with non-parametric tests or were rank-transformed before conducting parametric testing if they did not fulfill normality. Multiple comparisons were corrected for False Discovery Rate (FDR) to mitigate the likelihood of Type 1 error by employing the Benjamini-Hochberg procedure at 5% (Benjamini & Hochberg, 1995). All reported p-values were two-tailed.

Group differences in rank-transformed DDM parameters, SAT, accuracy, RT, and variance were assessed using independent t-tests. Tractography outcome measures (FA, MD, RD, AD, FR) and metabolite outcome measures (GABA, NAA, Glx, Myoinositol, Choline, Creatine) were compared between older and younger control groups by conducting non-parametric Mann Whitney U tests.

Hierarchical linear regression models were carried out for RT, SAT, and each EZ DDM parameter as dependent variables. Age and TOPF-UK score, the two variables that differed between the groups, were entered into the model first, followed by all metabolic and microstructural measurements in a stepwise fashion. Regression analyses were conducted on rank-ordered variables to account for non-normality of the data.

Following this, Pearson correlations on rank-transformed data were conducted between age, brain predictors and DDM parameters and the directionality of these relationships were explored with mediation analyses.

## 3. Results

### 3.1 Group differences in visual acuity

No significant differences were found in visual acuity between older and younger age groups (F(1,49)=1.239, p=.276) (Figure 3).

**Figure 3.**
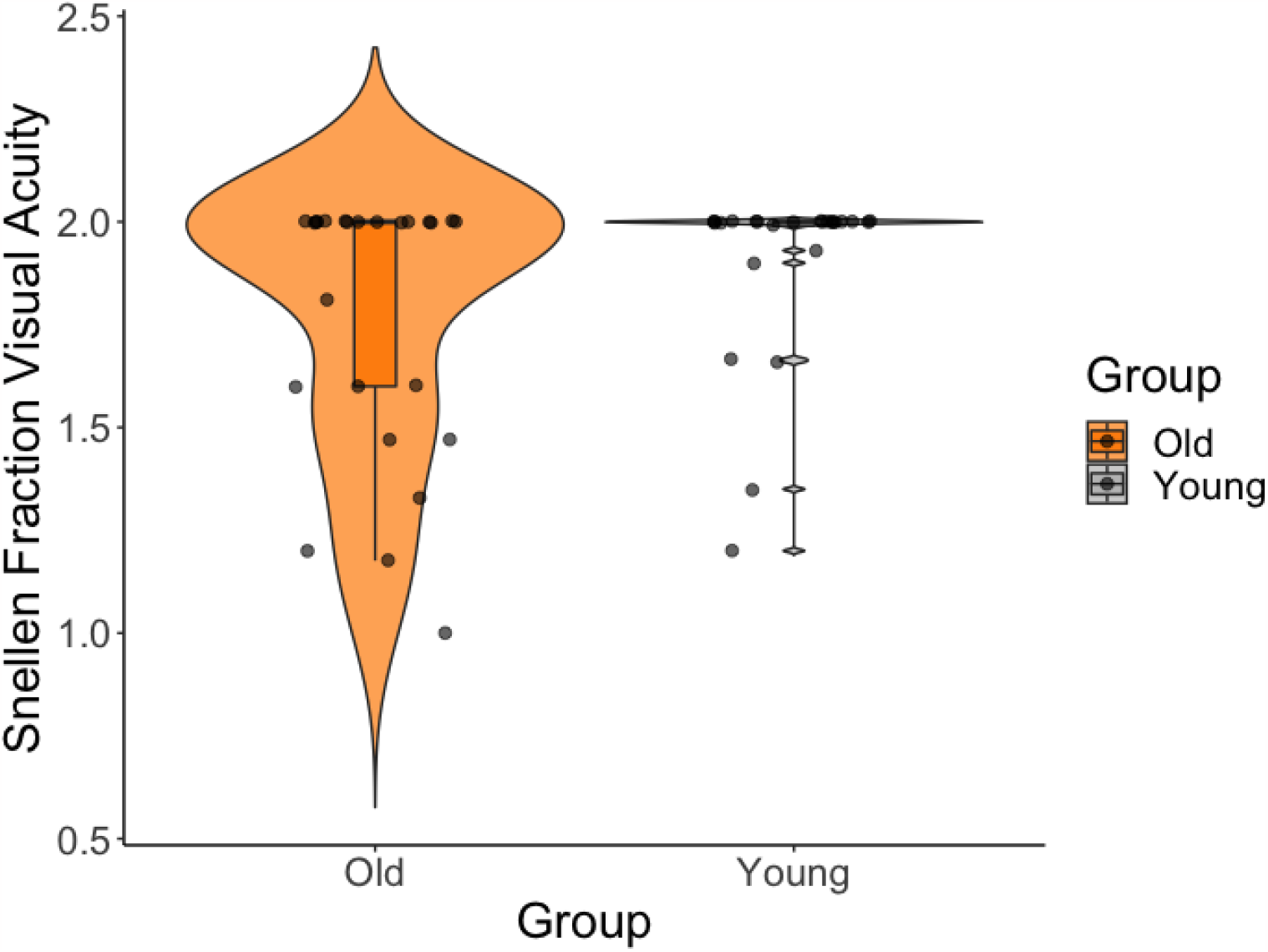
Violin plot with overlaid boxplot for group comparisons between older (orange) and younger (grey) adults’ visual acuity (Snellen Fraction). The boxplot displays the median and the interquartile range and the violin plot the kernel probability density, i.e., the width of the violin area represents the proportion of the data located there. There was no significant difference in visual acuity between younger and older adults.

### 3.2 Group differences in RT, SAT, and DDM parameters

Independent t-tests on rank-transformed data revealed that older compared to younger adults showed larger RT (t(48) = 4.2, p_FDRcor_ = 0.0007) (Figure 4C), non-decision time (t(48) = 2.9, p_FDRcor_ = 0.016) (Figure 4D) and boundary separation values (t(48) = 2.9, p_FDRcor_ = 0.016) (Figure 4E). No group differences were observed for accuracy (t(48) = 1.6, p = 0.12) (Figure 4A), SAT (t(48) = 0.18, p = 0.87) (Figure 4B) or drift rate (t(48) =0.16, p = 0.87) (Figure 4F).

**Figure 4.**
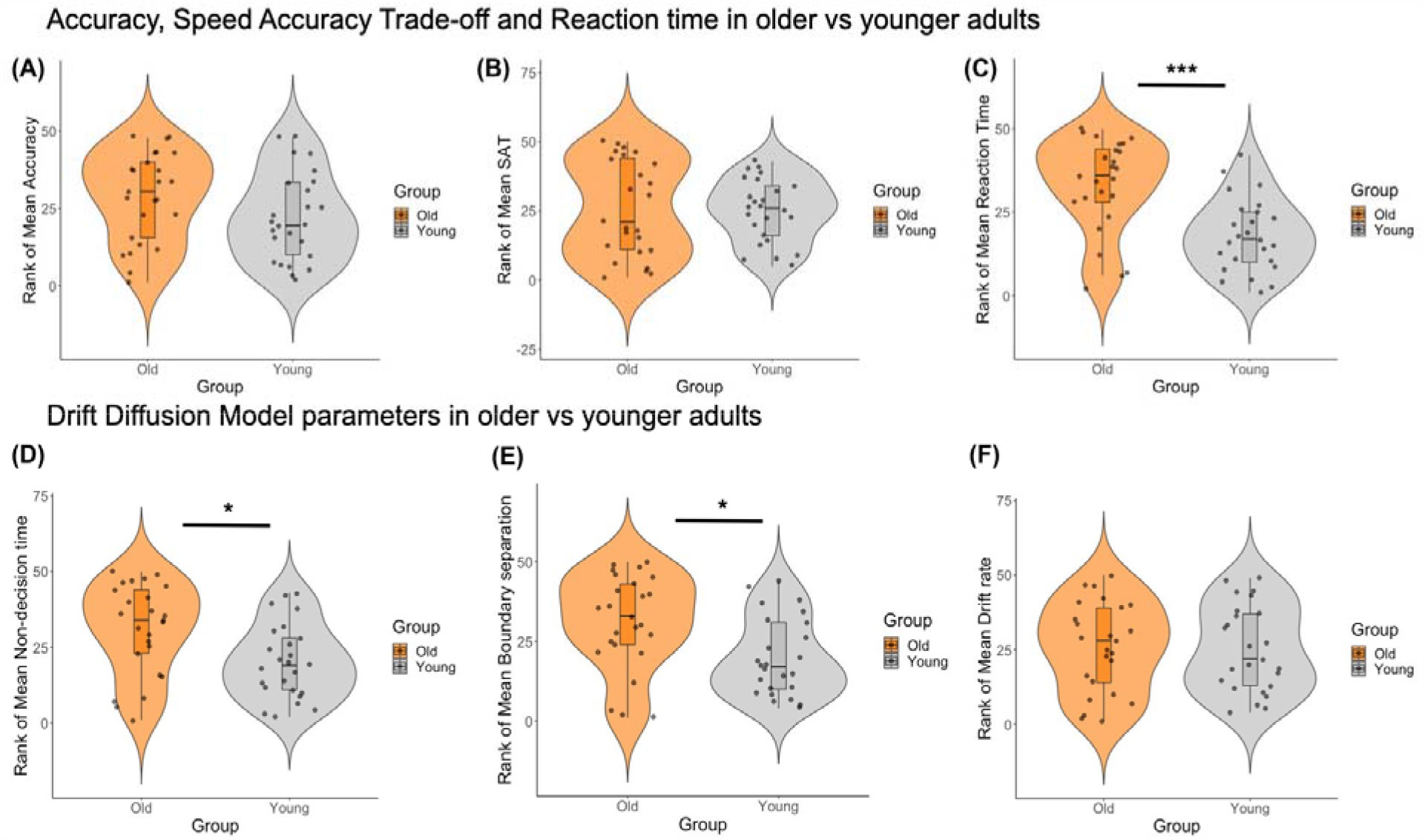
Violin plots with overlaid boxplots for group comparisons between older (orange) and younger (grey) adults’ rank-transformed accuracy, response time (RT), speed accuracy trade-off (SAT) performance and diffusion drift model (DDM) parameters. The boxplots display the median and the interquartile range and the violin plots the kernel probability density, i.e., the width of the violin area represents the proportion of the data located there. Older participants showed increased RT (mean rank-transformed RT_old_ = 33, SD =14.1; mean rank-transformed RT_young_ = 18, SD =10.9), boundary separation values (mean rank-transformed boundary separation_old_ = 31, SD =14.8; mean rank-transformed boundary separation_young_ = 20, SD =12.3) and non-decision time (perceptual and motor processing) (mean rank-transformed non-decision time_old_ = 30.9, SD =15; mean rank-transformed non-decision time_young_ = 20.1, SD =12.2). *** p_FDRcor_ <.001, * p_FDRcor_ <.05

### 3.3 MRI results

#### 3.3.1 Metabolic differences between older and younger adults

Group comparisons between older and younger participants showed no significant differences in GABA levels in the ACC, OCC or the PPC (Figure 5a). Older participants had significantly lower Glx (U=189, p=.004) and NAA (U=109, p<.001) in the ACC than younger adults. A trend towards significantly lower myoinositol in older adults in comparison to younger adults in the ACC was also observed (U=234, p=.058). In the PPC, older adults showed significantly lower NAA (U=190, p=.011), and a trend towards lower Glx (U=231, p=.054) than younger adults.

**Figure 5.**
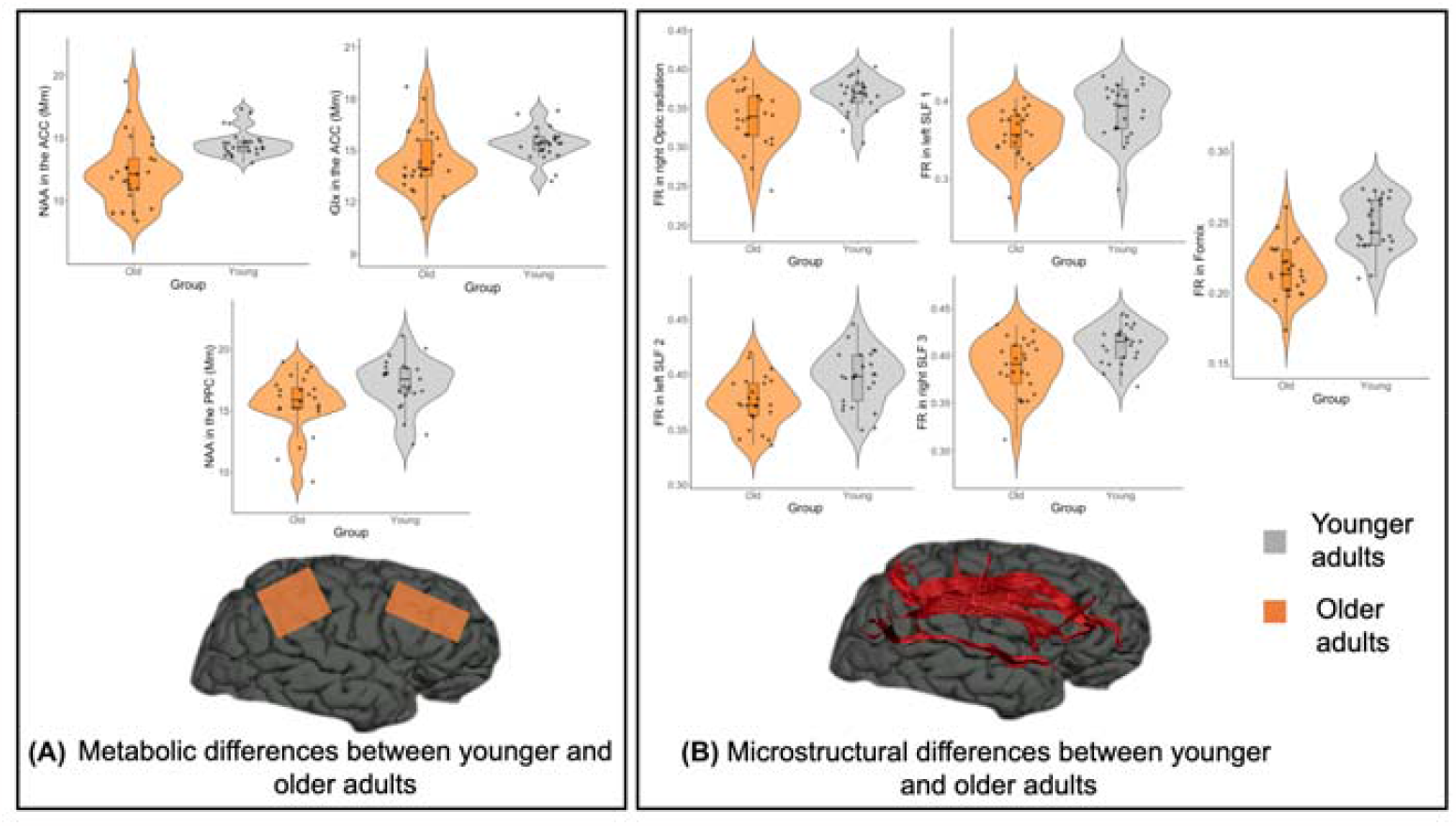
Metabolic and microstructural differences between younger and older adults. (A) Significant group comparisons for metabolites in voxels of interest between older and younger adults (B) Significant group comparisons for FR in tracts of interest between older and younger adults. FR was significantly lower in the fornix, optic radiation, SLF1, 2 and 3 in older adults (orange) in comparison to younger (grey) adults. **p<.001, *p<.0.05

#### 3.3.2 Diffusion weighted MRI differences between younger and older adults

Significantly lower restricted fraction was shown in the older group in the fornix (U=75, p<.001), right optic radiation (U=154, p=.001), left SLF1 (U=152, p=.001), left SLF2 (U=170, p=.002) and right SLF3 (U=169, p=.002) (Figure 5b).

Significantly higher FA in the fornix (U=22, p<.001), right optic radiation (U=207, p=.027), left ILF (U=203, p=.014), right ILF (U=157, p=.001), left SLF1 (U=131.5, p<.001), left SLF2 (U=195, p=.009), and right SLF2 (U=218, p=.029), and right SLF3 (U=163, p=.001) was found in younger adults in comparison to older adults. Significantly higher MD in the older group in comparison to the younger control group was found in the fornix (U=84, p<.001), left optic radiation (U=78, p<.001), right optic radiation (U=131, p<.001), right ILF (U=203, p=.014), right SLF1 (U=175, p=.003), right SLF3 (U=175, p=.003). Radial diffusivity was significantly higher in the older control group in all tracts of interest except the right SLF1, fornix (U=39, p<.001), left optic radiation (U=145, p=.001), right optic radiation (U=139, p<.001), left ILF (U=189, p=.007), right ILF (U=143, p<.001), left SLF1 (U=130, p<.001), left SLF2 (U=256, p=.199), right SLF2 (U=178, p=.003), left SLF3 (U=197.5, p=.010), right SLF3 (U=135.5, <.001) (Figure 6). Significantly greater axial diffusivity in older adults was found in the fornix (U=529, p<0.001), and significantly lower axial diffusivity in older adults was found in the SLF1 left (U=178, p=0.005).

**Figure 6.**
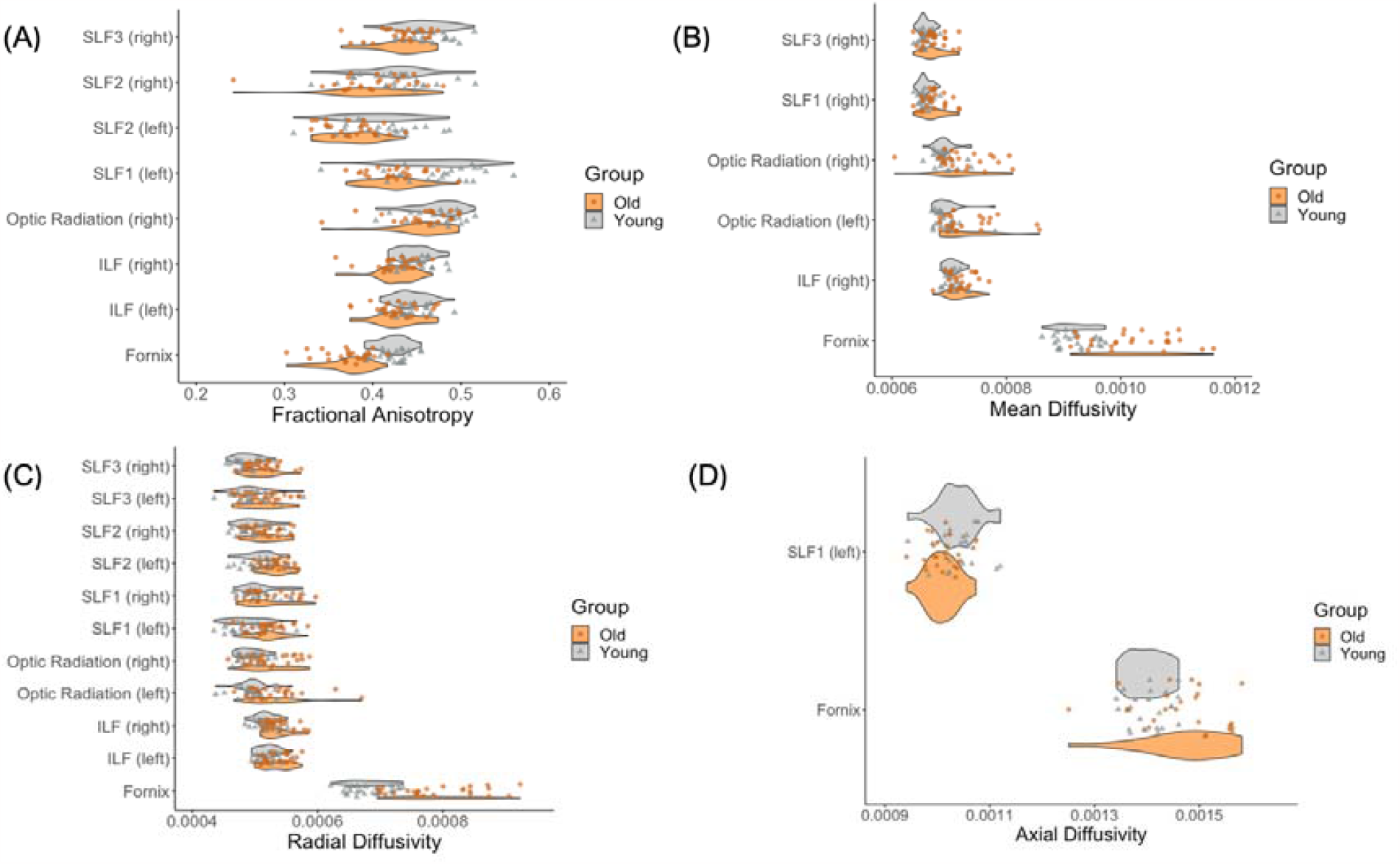
Diffusion tensor imaging (DTI) differences between younger and older adults. Significant group comparisons (p<.0.05) for tract fractional anisotropy (A), mean diffusivity (B), radial diffusivity (C) and axial diffusivity (D) between older and younger adults.

### 3.4 Correlations between response latency, SAT, and DDM parameters

Significant positive correlations were observed between mean RT and boundary separation (Rho = 0.74, p_FWEcor_ < 0.00000001), between SAT and boundary separation (Rho = 0.49, p_FWEcor_ = 0.002), and between mean RT and SAT (Rho = 0.4, p_FWEcor_ = 0.013). Drift rate and boundary separation were negatively correlated (Rho = -0.4, p_FWEcor_ = 0.013). Non-decision time did not correlate with any of the other cognitive variables (Figure 7).

**Figure 7.**
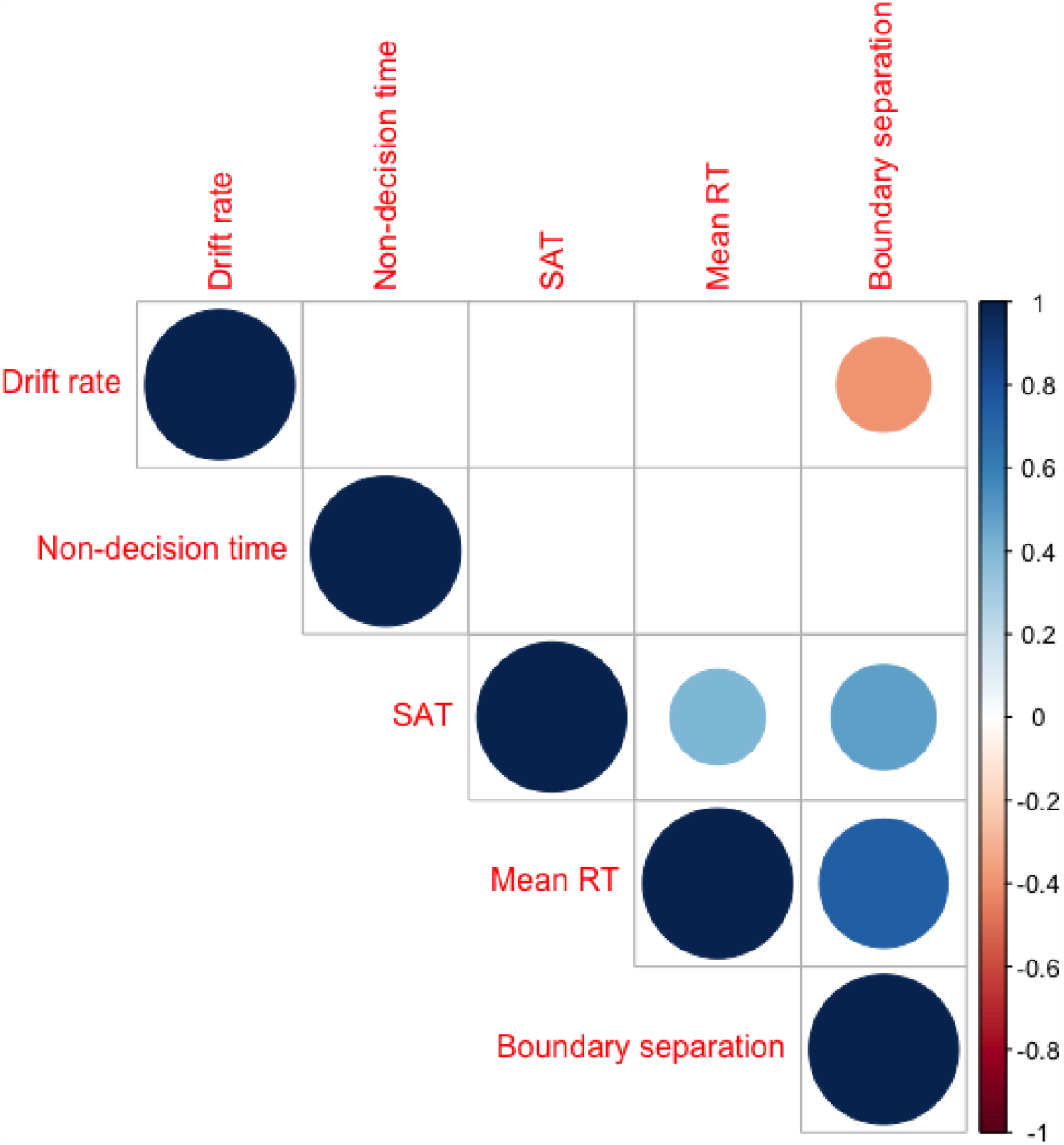
Correlation matrix between response latency, SAT and DDM parameters. Mean RT and boundary separation (p<.001), SAT and boundary sepataion (p<.001) and mean RT and SAT (p<.05) were positively correlated (blue shades). Drift rate and boundary separation were negatively correlated (p<.05) (red shade).

Mediation analysis revealed that boundary separation had a significant indirect effect (indirect ES of boundary separation = 0.31, SE =0.28, 95% CI 0.0005 - 0.985) and removed the direct effect of mean RT on SAT (remaining ES of mean RT on SAT = 0.09, SE = 0.18, 95% CI -0.289 – 0.471) (Figure 8). In contrast the inclusion of mean RT or SAT as mediator variables did not have any indirect effects on the direct effect of boundary separation on SAT (indirect ES of mean RT = 0.07, SE =0.24, 95% CI -0.51-0.35; direct ES of mean boundary separation on SAT = 0.42, SE = 0.18, 95% CI 0.043 – 0.8) or on mean RT (indirect ES of SAT = 0.027, SE =0.11, 95% CI -0.22 - 0.19; direct ES of mean boundary separation on mean RT = 0.71, SE = 0.11, 95% CI 0.49 – 0.94). This pattern of results demonstrates that differences in boundary separation accounted for the shared variance in mean RT and SAT.

**Figure 8.**
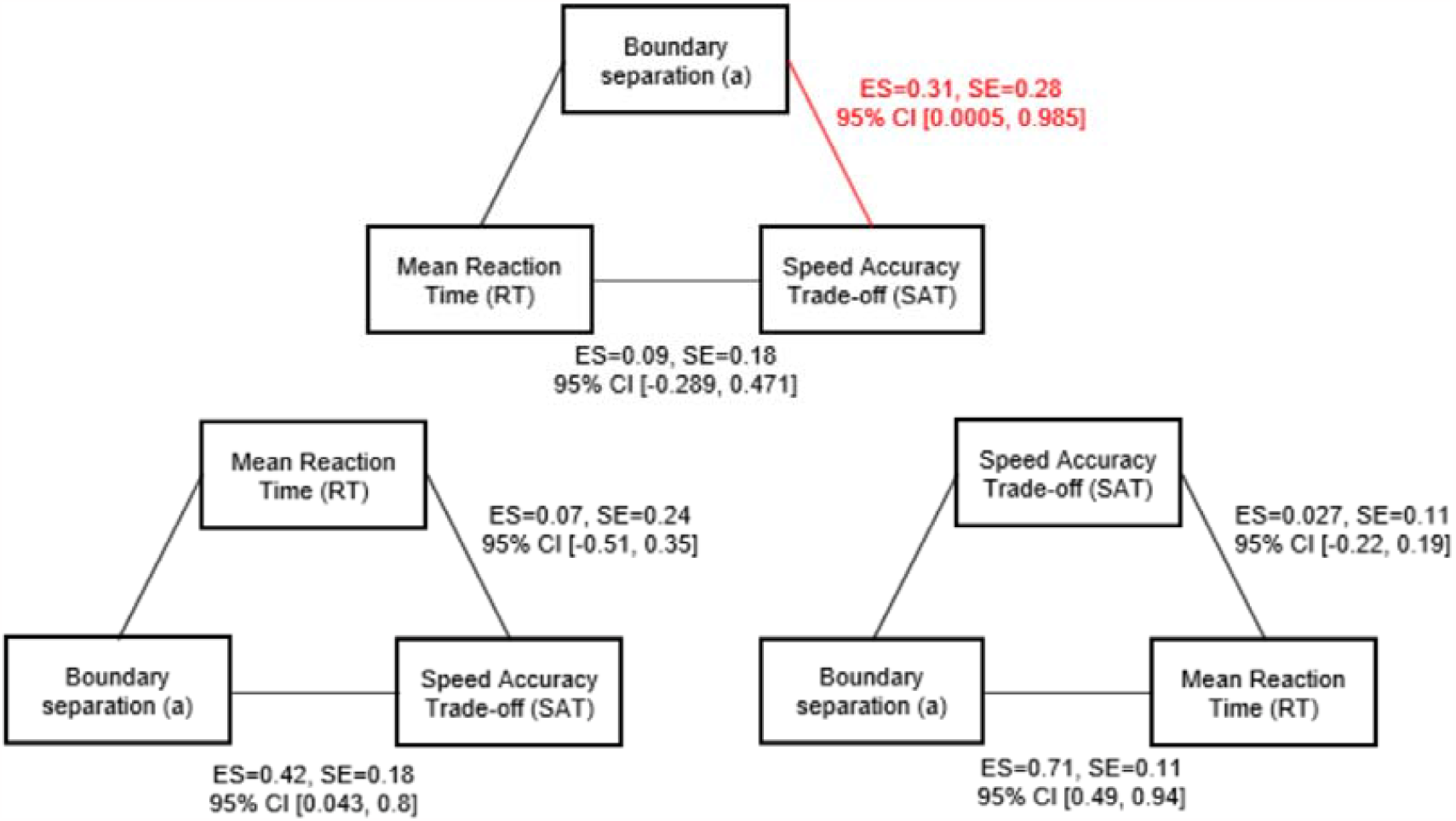
Mediation analyses between SAT, boundary separation and mean RT. Boundary separation had a significant indirect effect on SAT, removing the direct effect of mean RT on SAT. No other direct or indirect effects were significant.

### 3.5 Neurobiological predictors of response latencies, SAT, and DDM parameters

Hierarchical linear regression analyses testing for the effects of age, TOPF-UK score, and all microstructural and metabolic brain measurements on mean RT, SAT, and DDM indices were conducted separately for each outcome measure. All models entered age and TOPF-UK score as first predictors followed by the stepwise inclusion of the brain measurements.

#### 3.5.1 Response latencies (RTs) and SAT

Variation in RTs were not accounted for by age and TOPF-UK score along (adj R_2_ = 0.01, F(2,37) = 1.2, p=0.31) but the inclusion of the following microstructural and metabolic brain measurements improved the fit of the model significantly: fornix FA (delta R_2_=0.24, F(1,36)=12.5, p =0.001), AD in left optic radiation (delta R_2_=0.14, F(1,35)=8.9, p =0.005), RD in right SLF1 (delta R_2_=0.11, F(1,34)=8.7, p =0.006), myoinositol in OCC (delta R_2_=0.05, F(1,33)=4.2, p =0.048), and AD in right SLF1 (delta R_2_=0.06, F(1,32)=5.2, p =0.029). The final model explained 66% of the variation in RTs (adj R_2_ = 0.59, F(7,32) = 9.01, p<0.001) and included the following predictors: fornix FA (beta = -0.9, p_FDRcor_ < 0.0000001), RD in right SLF1 (beta = -0.33, p_FDRcor_ = 0.019), myoinositol in OCC (beta = 0.38, p_FDRcor_ = 0.019), TOPF-UK score (beta = 0.32, p_FDRcor_ = 0.025), AD in right SLF1 (beta = 0.35, p_FDRcor_ = 0.034) and AD in left optic radiation (beta = -0.26, p_FDRcor_ = 0.034).

Age and TOPF-UK score alone did not predict variability in SAT (adj R_2_ = -0.025, F(2,37) = 0.53, p=0.6). The inclusion of the following metabolic and microstructural measurements improved the fit of the model significantly (adj R_2_=0.42, F(5,34) = 4.9, p = 0.002): NAA in the ACC (delta R_2_=0.21, F(1,36)=10.12, p =0.003), RD in right SLF1 (delta R_2_=0. 1, F(1,35)=5.3, p =0.03) and FA in right SLF1 (delta R_2_=0.08, F(1,34)=4.5, p =0.04). In the final model SAT was significantly predicted by NAA in the ACC (beta = 0.5, p_FDRcor_ = 0.015) and RD in the right SLF1 (beta = -0.9, p_FDRcor_ = 0.015) (Figure 9).

**Figure 9.**
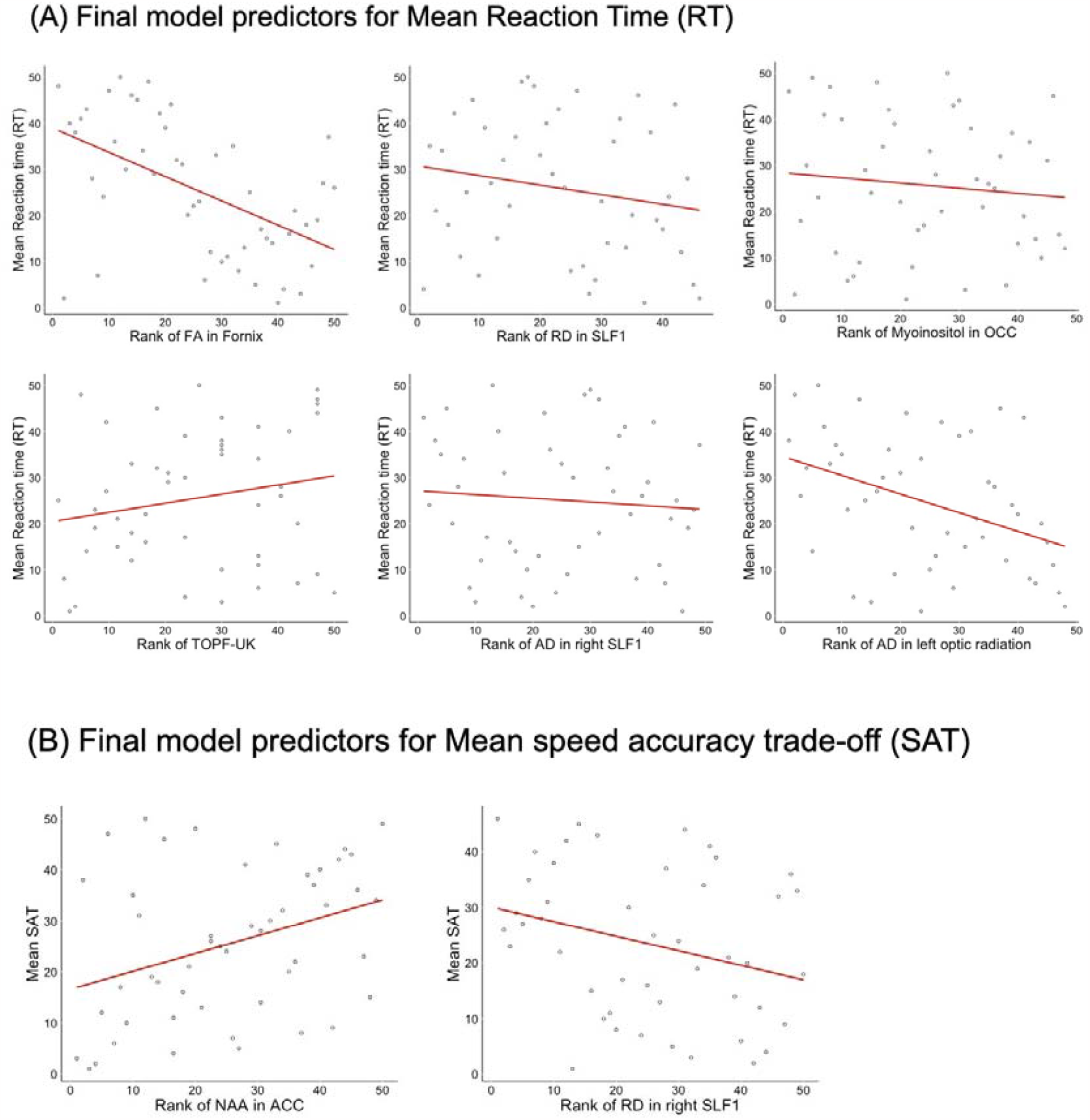
Slopes of the regression lines for the brain predictors in the final model for rank of mean reaction time (RT) (A) and mean SAT (B). Significant predictors (p<.05) of rank mean RT and rank mean SAT included in final hierarchical models.

#### 3.5.2 DDM parameters

Variation in boundary separation was not explained by age and TOPF-UK score (adj R_2_ = 0.029, F(2,37) = 1.6, p=0.22) but the inclusion of fornix FA (delta R_2_=0.20, F(1,36)=10.12, p =0.003) and FR in the right ILF (delta R_2_=0.08, F(1,35)=4.6, p =0.04) improved the fit of the model significantly (adj R_2_ = 0.29, F(4,35) = 5.01, p=0.003). In the final model fornix FA (beta = -0.8, p_FDRcor_ = 0.002) was the only significant predictor for variation in boundary separation.

Similarly, variation in non-decision time was not explained by age and TOPF-UK score alone (adj R_2_ = - 0.02, F(2,37), p=0.5). The inclusion of the following metabolic and microstructural measurements improved the model fit significantly (adj R_2_=0.47, F(6,33) = 6.7, p < 0.001: NAA in the ACC (delta R_2_=0.23, F(1,36)=11.5, p =0.002), AD in the right ILF (delta R_2_=0.15, F(1,35)=8.9, p =0.005), creatine in the OCC (delta R_2_=0.07, F(1,34)=4.3, p =0.045) and GLx in PPC (delta R_2_=0.07, F(1,33)=4.9, p =0.034). In the final model NAA in the ACC (beta = -0.45, p_FDRcor_ = 0.015), AD in the right ILF (beta = 0.38, p_FDRcor_ = 0.015), and creatine in the OCC (beta = 0.32, p_FDRcor_ = 0.03) predicted non-decision time significantly.

Finally, age and TOPF-UK score did not account for variation in drift rate (adj R_2_ = 0.01, F(2,37), p=0.32) and the inclusion of brain measurements did not improve the model fit significantly (adj R_2_=0.12, F(3,36) = 2.6, p = 0.07; delta R_2_=0.12, p=0.031) (Figure 10).

**Figure 10.**
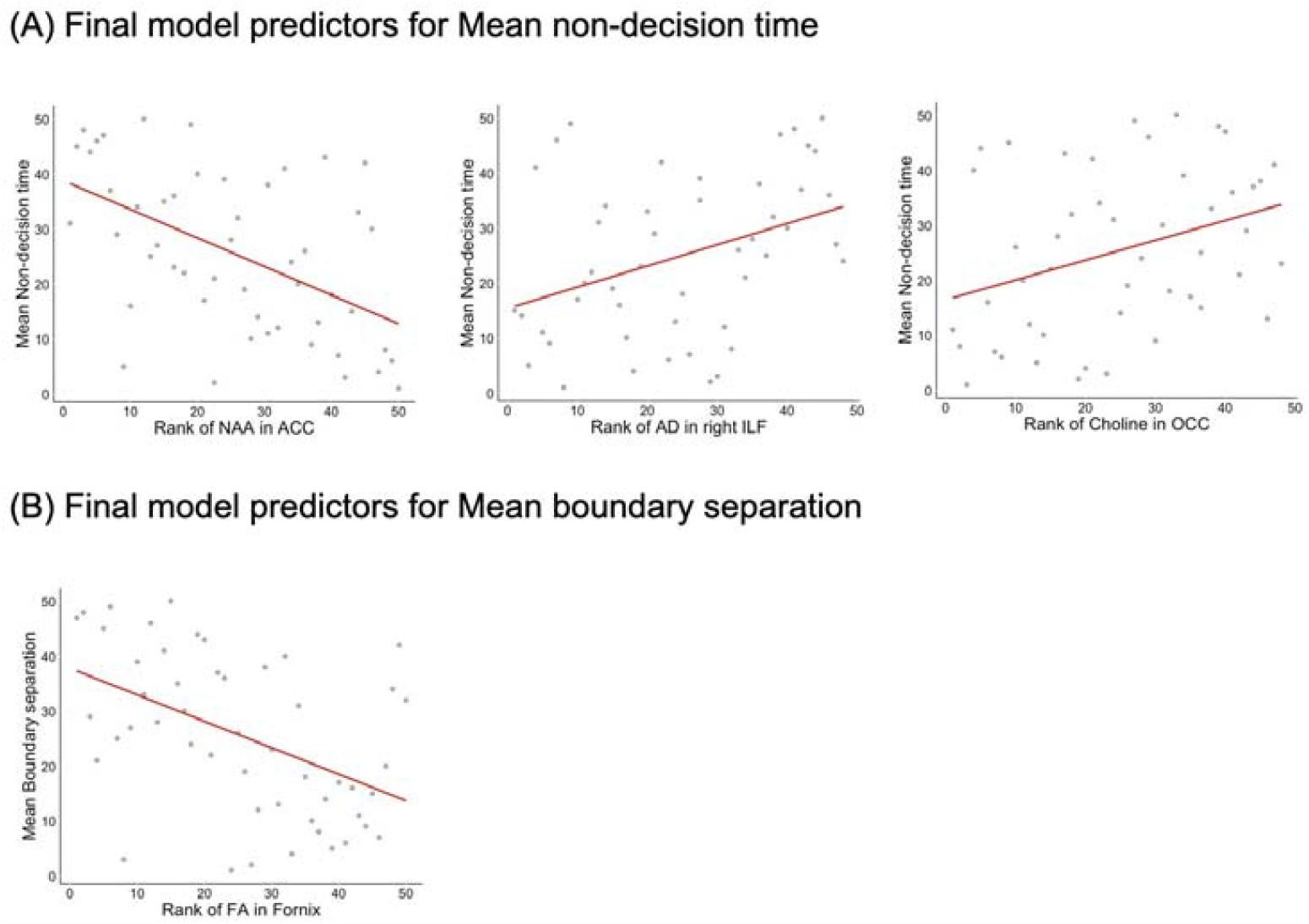
Slopes of the regression lines for the brain predictors in the final model for rank of mean non-decision time (A) and mean boundary separation (B). Significant predictors (p<.05) of rank mean non-decision time and rank mean boundary separation included in final hierarchical models.

#### 3.5.3 Correlation analyses between age, fornix FA, and RT

Spearman correlation coefficients were calculated between age, fornix FA, RT and boundary separation to explore the directionality of these relationships. Fornix FA correlated negatively with age (Rho = -0.75, p_FWEcor_ < 0.00000001), boundary separation (Rho = -0.48, p_FWEcor_ = 0.007), and RT (Rho = -0.53, p_FWEcor_ = 0.002). A significant positive correlation was present between age and RT (Rho = 0.35, p_FWEcor_ = 0.016) and a trend for a positive correlation between age and boundary separation (Rho = 0.26, p = 0.06). Mediation analysis demonstrated that fornix FA had a significant indirect effect (ES = 0.45, SE =0.13, 95% CI 0.21 - 0.7) and removed the direct effect of age on mean RT (remaining ES = -0.09, SE = 0.18, 95% CI -0.47 – 0.27). In contrast the inclusion of age as mediator variable did not have an indirect effect (ES = 0.07, SE =0.13, 95% CI -0.2 - 0.3) on the direct effect of fornix FA on RT (ES = -0.6, SE = 0.18, 95% CI -0.97 – -0.22).

This pattern of results suggests that age-related response slowing is mediated by the age-related decline in fornix microstructure but not vice versa (Figure 11).

**Figure 11.**
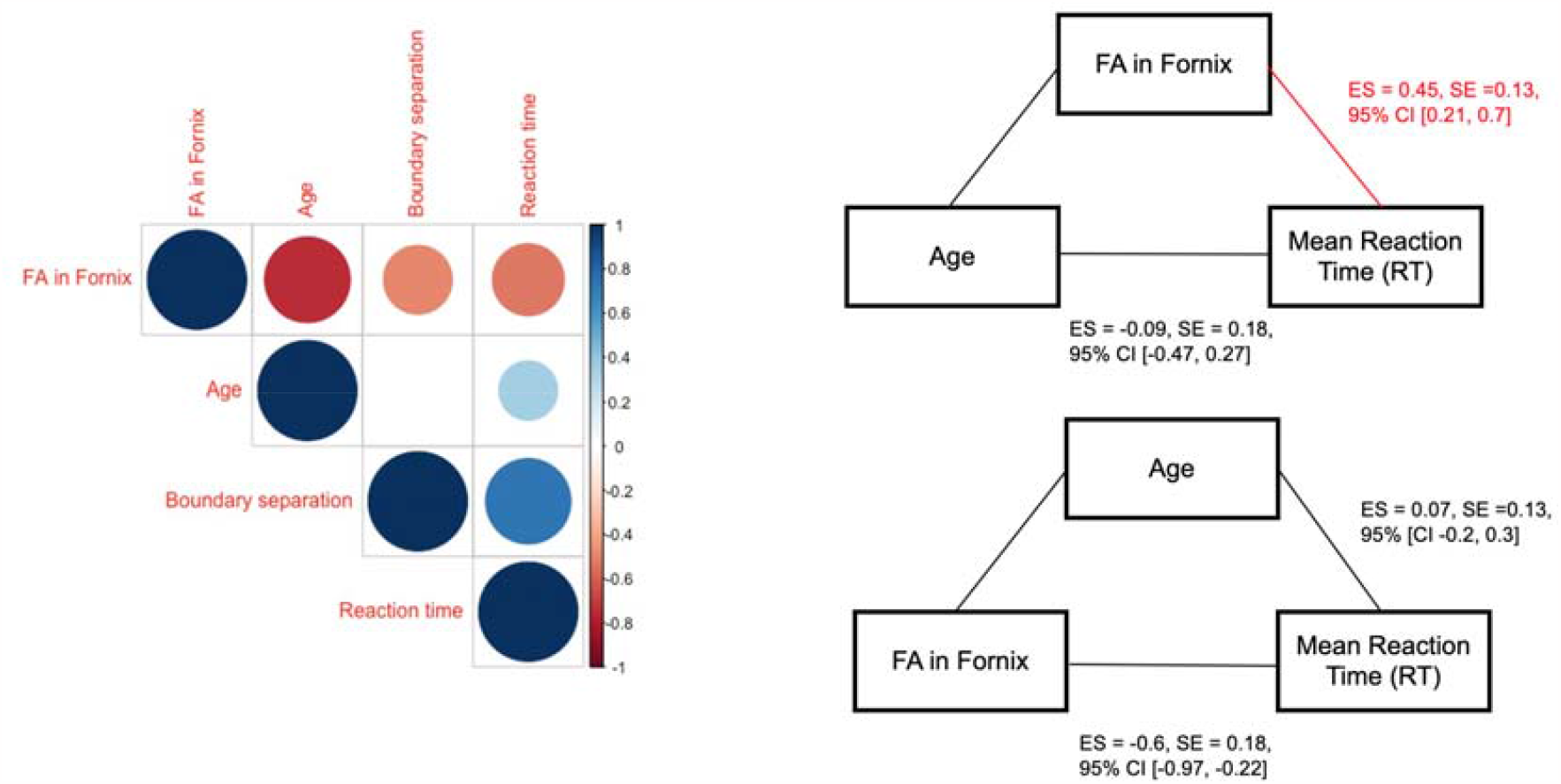
Correlation matrix and mediation analyses investigating the relationship between Age, FA in the Fornix, Mean Reaction time (RT) and boundary separation value. Correlation matrix (left) shows significant negative correlations (red share) (p<.05) between FA in fornix, age, mean RT and boundary separation. A significant positive correlation (blue shade) was present between age and RT. Mediation analyses (right) shows a significant indirect effect of FA in fornix on mean RT, which removed the direct effect of age on mean RT. Age as an indirect predictor did not remove the effect of FA in fornix on mean RT.

## 4. Discussion

The aim of this study was to investigate the cognitive and neurobiological substrates of age-related response slowing in visuo-perceptual decision making. For this purpose, we modelled older and younger adults’ RT data from the ANT flanker task with the EZ DDM to derive three cognitive components thought to contribute to response slowing in ageing. These were the non-decision time, that reflects the time needed for bottom-up sensory and top-down motor execution processes, the boundary separation value, that reflects the degree of conservatism regarding a response decision criterion, and the drift-rate, the speed with which information is accumulated to reach a response threshold. Furthermore, we employed advanced multi-shell diffusion-weighted MRI and MRS techniques to acquire estimates of micro-structural and metabolic properties of gray matter ROIs and white matter connections in visual-perceptual and attention networks. We then tested which of these brain measurements were predictive of differences in cognitive parameters for the purpose of elucidating the neurobiological basis of any age-related differences in visuo-perceptual decision making.

Consistent with the literature (Ratcliff et al., 2007; Rabbit, 1979; Theisen et al., 2021) and our hypotheses, we found age-related increases in RTs, boundary separation, and non-decision time. These differences were observed in participants without visual acuity impairments as measured with the Snellen Fraction, thus were not due to reduced eyesight in older participants. Contrary to our hypotheses, older adults did not show increased SAT, as measured with LISAS, and did not differ in accuracy or diffusion drift rates from younger adults.

Slowed response times while maintaining accuracy is a characteristic pattern observed in ageing and is generally thought to reflect a shift to a more conservative decision criterion that favours accuracy over speed. Consistent with this view we observed age-related increases in boundary separation. Importantly, mediation analysis revealed that the differences in boundary separation accounted for the shared variance between SAT and RTs but not *vice versa*, suggesting that the shift to a more conservative response strategy is the main contributor to the slowing of decision-making in older age. Drift rate was negatively correlated with boundary separation reflecting that a more conservative response strategy involves the accumulation of more information, and hence reduces the speed of reaching a response threshold criterion. Non-decision time did not correlate with any of the other cognitive variables, suggesting that this variable captures other processes than those involved in central decision-making. This result accords with the DDM model’s assumption that non-decision time comprises low-level sensory and pure motor execution processes that are independent of central decision-making processes, such as boundary separation. Thus, although ageing increased non-decision time, as proposed by the sensory degradation and the motor noise hypotheses, this age-related decline in sensory-motor function was unrelated to the increase in boundary separation and only the latter accounted for overall response slowing. In summary, the pattern of results for the cognitive data indicates that older adults responded more slowly because they adopted a more conservative criterion threshold that favoured responding accurately over responding quickly (Rabbitt, 1979; Salthouse, 1979; Smith & Brewer, 1995; Starns & Ratcliff, 2010). While age-related increases in non-decision time were observed and accord with accounts of low-level sensory degradation and slowing of motor execution with age, these processes did not contribute to overall response slowing or to the increases in boundary separation.

The main aim of our study was to investigate the neurobiological basis of differences in cognitive components thought to underpin age-related response slowing. For this purpose, we acquired microstructural and metabolic measurements with advanced diffusion-weighted MRI and MRS from key regions within visual perceptual and attention-executive networks. Regions involved in bottom-up visuo-sensory and perceptual processing comprised the occipital cortex as well as the optic radiation and ILF while selected areas hypothesized to mediate top-down decision-making processes were the ACC, PPC, SLF and fornix. Consistent with our hypotheses and the literature (Michielse et al., 2010; Salat et al 2005; Yang, Bender & Raz, 2015), we observed age-related reductions in FA and FR and increases in MD, RD and AD indices in all white matter pathways, demonstrating that they were detrimentally affected by age. In contrast to previous studies, we employed not only DTI measurements but also the restricted signal fraction FR from CHARMED. DTI measures are not only affected by the biological properties of white matter such as myelin and axon density but also by the geometry and complexity of fiber architecture for instance crossing fibers, and hence are difficult to interpret in terms of any specific tissue changes (De Santis et al., 2014). FR from CHARMED reflects the fraction of the signal from a restricted diffusion compartment that is thought to arise from intra-axonal space in white matter. Reductions of FR have been proposed to reflect a decrease in the density of axons, that may occur due to a loss of myelin and/or axons secondary to Wallerian degeneration in ageing (De Santis et al., 2014).

Older relative to younger adults also showed reduced levels of NAA and Glx in the ACC and lower levels of NAA in the PPC. NAA and Glx are both markers of neuronal metabolism (Newsholme et al., 2003) and are considered to play a key role in energy metabolism in neural mitochondria (Lu et al., 2004). These findings are consistent with accumulating evidence of energy depletion as a key component of biological aging (Raz & Daugherty 2018). With normal aging, the accumulation of biological ‘imperfections’ such as protein aggregation are thought to impair mitochondrial function and cause low-level inflammation, resulting in reduced glucose uptake, synaptic deterioration, and gliosis (Currais, 2015), and in turn to further energy reduction. The here observed reductions in NAA and Glx in the ACC in older adults may reflect a decreased neuronal metabolism due to the accumulation of these biological ‘imperfections’. The pattern of our results suggest that aging affects neuronal metabolism primarily in frontal and parietal attention regions but not in the OCC, consistent with evidence suggesting that healthy aging is particularly associated with a reduction in mitochondrial energy metabolism in frontal brain regions (Reddy & Beal, 2008, Reutzel et al., 2020, Yin et al., 2014).

Hierarchical regression analyses testing for effects of the microstructural and metabolic brain measurements on DDM components while controlling for age and TOPF-UK score demonstrated that differences in boundary separation were predicted by differences in fornix FA, while differences in non-decision time were predicted by regions involved in both bottom-up and top-down processes, specifically NAA in the ACC, AD in the right ILF, and creatine in the OCC. No brain predictors were found for drift rate. In addition, differences in fornix FA together with diffusivities (AD, RD) in right SLF1, myoinositol in OCC, AD in left optic radiation, and the verbal intelligence as estimated by the TOPF-UK score predicted differences in overall RT while differences in NAA in the ACC and RD in the right SLF1 predicted differences in SAT. Age was not a significant predictor in any model.

The observed pattern of relationships between brain measurements and cognitive components were overall consistent with our hypotheses. More specifically we observed that differences in microstructural and metabolic tissue properties in top-down regions known to be involved in decision making (fornix, ACC, SLF) predicted boundary separation (fornix) and SAT (ACC, SLF) while a combination of measurements from bottom-up sensory (OCC, ILF) and top-down motor execution (ACC) regions predicted non-decision time that captured low level sensory function and pure motor execution. Finally, differences in overall response time were predicted by brain estimates from a network of top-down (fornix, SLF) and bottom-up brain regions (OCC, optic radiation) together with verbal intelligence, reflecting that response latency is a function of all of the processes involved in visuo-perceptual decision-making.

Fornix FA was the only brain predictor for boundary separation and the largest predictor for overall response time while age did not contribute to the regression models. The fornix is the main output connection from the hippocampus to other limbic and cortical regions and is known to be detrimentally affected by age (Chen et al., 2015; Metzler-Baddeley et al 2011). The role of the hippocampus and the fornix in episodic memory processing is well-established and age-related decline in fornix microstructure has been shown to contribute to episodic memory impairments in ageing (Douet & Chang, 2015; Metzler-Baddeley et al 2011). However, based on neuroimaging and lesion studies that have shown an involvement of the hippocampus and the fornix in complex visual discrimination tasks (Lech et al 2016; Postans et al., 2014), it has been proposed that the medial temporal lobe structures do not only process mnemonic but also visuo-perceptual and spatial functions (Graham et al 2010; Lech et al 2016).

To better understand the role of the fornix in age-related slowing of visual-perceptual decision making, we carried out additional correlational analyses that revealed large negative correlations between fornix FA and age (Rho = -0.75) as well as between fornix FA and RT (Rho = -0.5) and fornix FA and boundary separation (Rho = -0.48 demonstrating that lower fornix FA values were associated with slower response times and higher boundary separation values. Finally, aging was positively correlated with RT (Rho = 0.35) and a trend for a positive correlation between age and boundary separation (Rho = 0.26) was observed. Additional mediation analyses revealed that differences in fornix FA removed any effects of age on RT while the inclusion of age did not remove the effects of fornix FA on RT and boundary separation. This pattern of results indicates that age-related response slowing was mediated by the agerelated decline in fornix microstructure but not *vice versa*. In other words, fornix FA mediated the relationship between age and increase in response time. Fornix FA also predicted boundary separation, which fully explained the shared variance in SAT and RT.

To summarise, our findings indicate that age-related slowing in visual discrimination is primarily driven by the adoption of a more conservative response strategy. The degree of conservatism adopted relies on fornix microstructure, regardless of age. However, as fornix microstructure declines with advancing age possibly due to axonal loss and/or myelin damage, age is indirectly associated with the adoption of a more conservative decision strategy. Based on this pattern of results we propose that the fornix plays an important role in the accumulation of information to reach a response threshold criterion in a visual perceptual discrimination task. If fornix integrity is compromised, for instance by ageing, this process will become noisier and more information needs to be accumulated before a decision can be reached leading to a more conservative response strategy.

We also observed an increase of non-decision time with age, which was predicted by a reduction of NAA in the ACC and increases of creatine in the OCC and AD in the right ILF. This suggest that age-related decline in visual sensory-perceptual networks and regions involved in motor action control (ACC) due to reduced energy metabolism and neural activity, myelin damage and/or axonal loss underpin loss of sensory motor functions. While the differences in brain regions involved in sensori-motor functions predicted overall response slowing and non-decision time, non-decision time was unrelated to differences in fornix microstructure, boundary separation and overall RT. Thus, we did not find any evidence to support the view that age-related sensory motor decline in terms of sensory degradation and motor noise, led to the adoption of a more conservative response strategy.

A few limitations of our study are noteworthy. Firstly, the sample size of 25 participants per age groups was relatively small and it would be advantagous to replicate the here observed findings in a larger cohort. However, we did successfully replicate a number of well-established effects notably the increase of non-decision time, boundary separation, and overall RT, the widespread reductions of white matter micro-structure and of ACC and Glx in older versus younger adults. Our main findings are also based on moderate to large effect sizes, suggesting that despite the modest sample size we were appropriately powered to study the cognitive and neurobiological correlates of age-related slowing in visuo-perception.

Our results suggest that under the specific task conditions employed here boundary separation may provide a more sensitive measurement of SAT than LISAS for which no age effect was present. Our instructions for the ANT flanker task did not seek to experimentally manipulate accuracy-speed trade-offs by emphasising speed over accuracy or *vice versa*. It could therefore be that opposing between-subject trade-offs may have masked any SAT differences between the age groups. In other words, the increased speed by one participant may have been compensated by the increased accuracy by another. LISAS is a linear measure of SAT that has been shown to be insensitive to SAT, if the SAT effects are linearly balanced across participants (Vandierendock 2017). Thus, LISAS may have been a suboptimal choice to estimate SAT in the present study.

Finally, the neurobiological interpretation of the differences in white matter microstructural indices needs to be done with caution. While we observed age-related reductions in FR from CHARMED in the fornix, optic radiation and SLF, suggesting a decline in axon density and/or axonal myelination in these pathways, the FR index did not significantly predict any cognitive components. In contrast, the DTI measurements of FA, axial and radial diffusivities were identified as significant brain predictors of cognition in the regression analyses. This pattern of results suggests that DTI indices that capture a variety of age-related differences in white matter microstructure due to differences in biological properties, such as loss of myelin and axons and increased inflammation, and differences in the geometry and complexity of fibers are more sensitive brain predictors of cognitive change than the FR index that is thought to be more sensitive to specific biological properties of white matter (axon density). DTI and FR measures were corrected for partial volume effects with the Free Water Elimination Method (Pasternak et al., 2009), but this method cannot completely rule out that atrophy-induced free water contamination may have biased these indices. This may have been particularly the case for DTI indices notably in regions susceptible to partial volume contamination such as the fornix (Metzler-Baddeley et al 2012; see Parker et al., 2021) and may have made these indices more sensitive to age-related tissue atrophy.

### 4.2 Conclusions

Our study provides novel insights into the cognitive and neurobiological underpinnings of age-related response slowing. We demonstrate that age-related response slowing occurs due to the adoption of a conservative response strategy that arises from the decline in fornix microstructure that accompanies aging rather than reflecting a direct effect of age. We further provide evidence that age-related increases in response times and boundary separation are unrelated to increases in non-decision time due to impaired sensory-motor functions. Finally, we identified that age-related reductions in metabolic and microstructural properties of gray and white matter regions within bottom-up and top-down visual perceptual decision making networks.

