## Supplementary tables for "Age-related fornix decline predicts conservative response strategy-based slowing in perceptual decision-making"

Supplementary Table 1: Enter and stepwise regression model outcomes for predictors of SAT, mean RT and DDM parameters

| **Speed accuracy trade-off (SAT)** | | | | | |
| --- | --- | --- | --- | --- | --- |
| **Predictors** | **R2** | **Adj R2** | **F test** | **Beta** | **significance** |
| TOPF |  |  |  | 0.208 | t=1.213, p=0.234 |
| Age |  |  |  | 0.013 | t= 0.089, p=0.930 |
| NAA in ACC |  |  |  | 0.479 | t= 2.967, p=0.005 |
| RD in SLF1 (right) |  |  |  | -0.986 | t= -2.951, p=0.006 |
| FA in SLF1 (right) |  |  |  | -0.710 | t= -2.2126, p=0.041 |
| Final model:  TOPF, Age, NAA in ACC, RD in SLF1 (right), FA in SLF1 (right) | 0.419 | 0.333 | F(5,34)=4.896, p=.002 |  |  |
| **Mean Reaction time (RT)** | | | | | |
| **Predictors** | **R2** | **Adj R2** | **F test** | **Beta** | **significance** |
| TOPF |  |  |  | -0.058 | t=-0.305, p=0.763 |
| Age |  |  |  | 0.322 | t=2.596, p=0.014 |
| Fornix FA |  |  |  | -0.911 | t=-5.253, p<.001 |
| AD in optic radiation (left) |  |  |  | -0.263 | t=-2.288, p=0.029 |
| RD in SLF1 (right) |  |  |  | -0.330 | t=-2.953, p=.006 |
| Myoinositol in OCC |  |  |  | 0.380 | t=2.844, p=.008 |
| AD in SLF1 (right) |  |  |  | 0.345 | t=2.284, p=.029 |
| Final model:  TOPF, Age, Fornix FA, AD in optic radiation (left), RD in SLF1 (right), Myoinositol in OCC, AD in SLF1 (right) | 0.663 | 0.590 | F(7, 32)=9.007, p=<.001 |  |  |
| **Mean Non-decision time (t)** | | | | | |
| **Predictors** | **R2** | **Adj R2** | **F test** | **Beta** | **significance** |
| TOPF |  |  |  | -0.131 | t=-0.879, p=0.386 |
| Age |  |  |  | 0.039 | t=0.292, p=0.772 |
| NAA in ACC |  |  |  | -0.453 | t=-3.109, p=.004 |
| AD in ILF (right) |  |  |  | 0.384 | t=2.979, p=.005 |
| Choline in OCC |  |  |  | 0.320 | t=2.589, p=0.014 |
| Glx in PPC |  |  |  | -0.273 | t=-2.215, p=0.034 |
| Final model: TOPF, Age, NAA in ACC, AD in ILF (right), Choline in OCC, Glx in PPC | 0.550 | 0.468 | F(6,33)=6.714, p<.001 |  |  |
| **Mean Boundary separation (a)** | | | | | |
| **Predictors** | **R2** | **Adj R2** | **F test** | **Beta** | **significance** |
| TOPF |  |  |  | -0.155 | t=-0.757, p=0.454 |
| Age |  |  |  | -0.180 | t=-1.195, p=0.240 |
| Fornix FA |  |  |  | -0.807 | t=-3.881, p<.001 |
| FR in ILF (right) |  |  |  | 0.348 | t=2.137, p=0.04 |
| Final model: TOPF, Age, Fornix FA, FR in ILF (right) | 0.364 | 0.292 | F(4,35)-5.013, p=.003 |  |  |
| **Mean Drift rate (v)** | | | | | |
| **Predictors** | **R2** | **Adj R2** | **F test** | **Beta** | **significance** |
| TOPF |  |  |  | 0.411 | t=2.364, p=0.024 |
| Age |  |  |  | -0.114 | t=-0.708, p=.483 |
| FA in ILF (right) |  |  |  | 0372 | t=2.262, p=.030 |
| Final model: TOPF, Age, FA in ILF (right) | 0.177 | 0.108 | F(3,36)=2.579, p=.069 |  |  |
